# SMART: an approach for accurate formula assignment in spatially-resolved metabolomics

**DOI:** 10.1101/2025.03.12.642824

**Authors:** Yinghao Cao, Shengxi Li, Zhaoyi Liu, Zixian Jia, Weidong Zhuang, Xu Pan, Jinyu Zhou, Lifeng Yang, Lin Wang

## Abstract

Spatially-resolved metabolomics plays a critical role in unraveling tissue-specific metabolic complexities. Despite its significance, this profound technology generates thousands of features, yet accurate annotation significantly lags behind LC-MS-based approaches. To bridge this gap, we introduce SMART, an open-source platform designed for precise formula assignment in mass spectrometry imaging. SMART constructs a KnownSet database containing 2.8 million formulas linked by DBEdges derived from repositories such as HMDB, ChEMBL, PubChem, and BioEdges from KEGG biological reactant pairs. Using a multiple linear regression model, SMART extracts formula networks associated with the *m/z* of interest and scores potential candidates based on several criteria, including linked formulas, DBEdges/BioEdges, and ppm values. Benchmarking against reference datasets demonstrates that SMART achieves prediction accuracy rates of up to 92.4%. Applied to mass spectrometry imaging, SMART successfully annotated 986 formulas in developing mouse embryos. This robust platform enables systematic formula annotation within tissues, enhancing our understanding of metabolic heterogeneity.

## Introduction

Metabolomics is a powerful tool for mapping the landscape of metabolites in biological systems, offering insights into cellular functions, disease progression, drug responses^1-3^, and bridging the genotype-phenotype gap^4, 5^. Liquid chromatography-Mass spectrometry (LC-MS)-based metabolomics allows simultaneous detection and quantification of thousands of metabolites features and is widely used in plants, microbial and mammalian studies. Advances in tools like XCMS^6^, MetLin^7^, MetDNA^8^, NetID^9^, SIRIUS^10^, BUDDY^11^ and KGMN^12^ have enabled the discovery of numerous biologically active metabolites.

Recent progress in mass spectrometry imaging (MSI) have propelled spatially-resolved metabolomics to the forefront, allowing visualization of metabolic processes within biological systems^13-21^ and probing metabolic fluxes, such as TCA carbon flux^22^, amino nitrogen flux^23^, and lipogenic flux^24^. However, challenges persist in untargeted spatially-resolved metabolomics, due to the chemical diversity and abundance variability of metabolites, and limitations like the lack of chromatographic separation, retention time, and sufficient ion abundance for robust MS/MS spectra. Although ion mobility and post ionization techniques offer potential solutions, high cost and complexity limit their broad application^25^. Consequently, LC-MS or LC-MS/MS-based metabolomics tools cannot be seamlessly adapted for MSI data analysis.

For metabolites annotation in MSI, formula assignment typically starts by matching *m/z* signals to metabolomic database like HMDB^26^ and KEGG^27^, or by identifying isotopic patterns based on ion intensity and spatial localization pattern. Multiple tools have been developed to improve confidence in formula annotation. For instance, pySM employes a false discovery rate (FDR)-controlled framework for metabolite annotation^28^, rMSIannotation integrates isotopic pattern coherence, image correlation and mass error^29^, and MSIannotator compares MSI data against reference metabolites from LC-MS data^30^. However, these methods have limitation in providing reliable annotations for MSI data due to two potential flaws: i) many compounds are either poorly characterized or absent in current databases, hindering accurate formula assignment for detected *m/z* values. ii) instrumental errors, along with limitations in mass accuracy, resolution and sensitivity, introduce uncertainties that further challenge annotation reliability.

To address these issues, we introduce SMART (Spatial Metabolite Formula Predictor, Figure 1a), a novel tool designed for precise formula annotation in MSI. SMART builds a comprehensive KnownSet formula database by integrating formulas from repositories including HMDB, ChEMBL^31^, and PubChem^32^. By analyzing chemical shifts (DBEdges) from random metabolite pairs in Knownset (Supplementary Table 1) and biological formula shift (BioEdges) from the KEGG REACTION^27^ database, SMART connects all possible formulas in KnownSet database by DBEdges/BioEdges. Based on 118 manually validated metabolites from NetID, SMART extracted several features from KnownSet database to create a multiple linear regression (MLR) model for formula annotation. Validation on large datasets shows that SMART achieves high prediction accuracy, reaching up to 92.4%. Applying SMART to MSI data enabled systemic annotation of 986 formulas in developing embryos, providing new insights into spatial metabolic compartmentalization and its dynamic alteration during embryogenesis.

**Fig 1.**
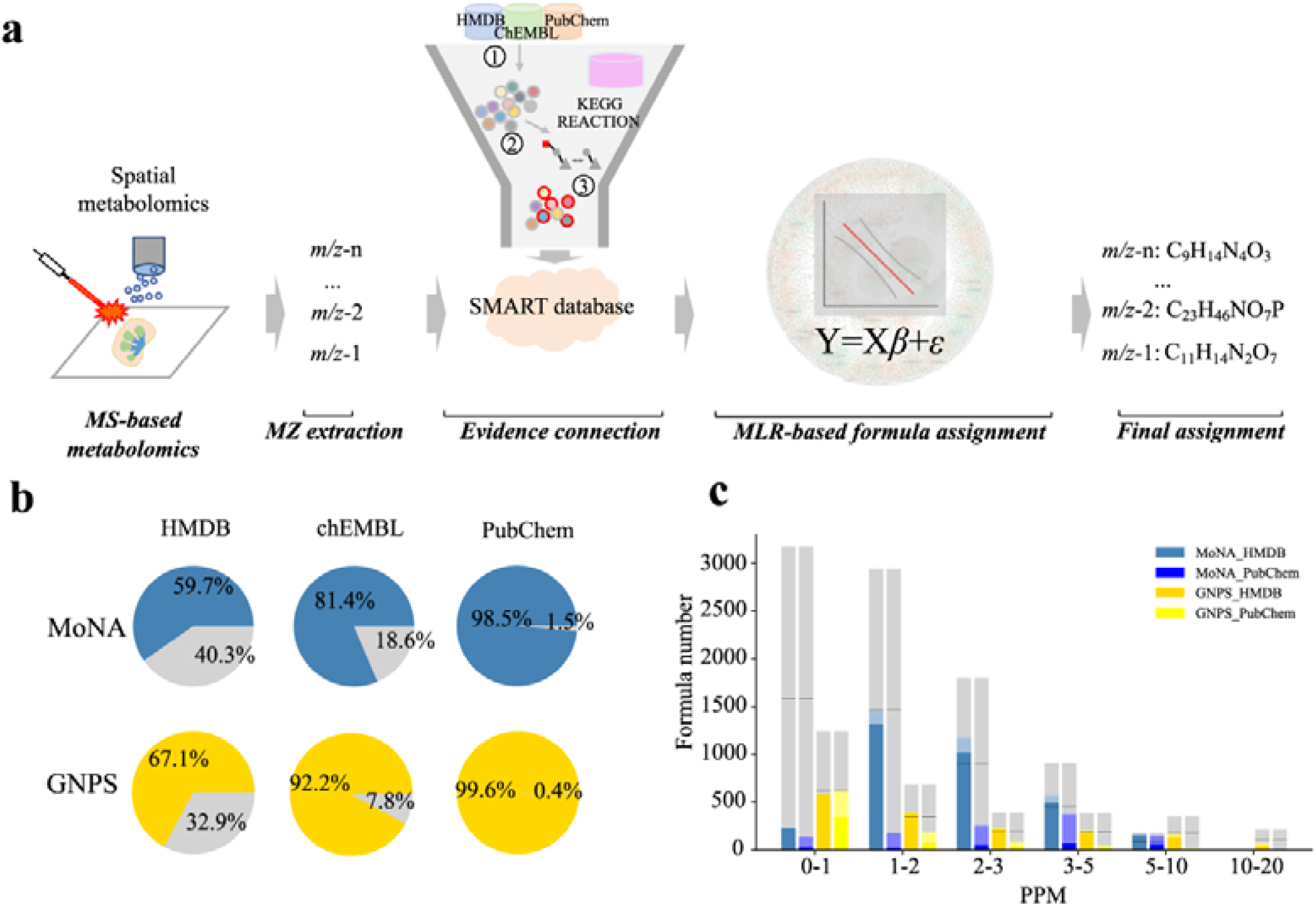
The framework of SMART and ppm matching for formula assignment. a. The framework of SMART. b. The formulas from MoNA and GNPS datasets existed in public databases. Gray color represents missing formulas. c. Formulas assignment for MoNA and GNPS datasets based on ppm matching against HMDB or PubChem. Gray color represents false assignment. For each bar, the dark color represents Top 1 accuracy, while the light color represents Top 3 accuracy, respectively. Black lines indicate the boundary for a prediction accuracy of 50%.

## Result

## Evaluation of Metabolite Formula Matching Across Databases Using ppm

HMDB and PubChem are prominent databases for metabolite matching and novel compound searches. To assess their coverage, we analyzed two datasets: MoNA (9,010 metabolites) .(https://mona.fiehnlab.ucdavis.edu/) and GNPS^33^ (3,280 metabolites) We compared their unique formulas with those in HMDB, PubChem and ChEMBL. Results showed that 40.3% of MoNA and 32.9% of GNPS formulas were missing in HMDB, while PubChem covered most formulas with only 0.4%-1.5% missing (Figure 1b). Next, we extracted the measured *m/z* values from MoNA and GNPS, grouping them into six ppm error intervals: 0-1, 1-2, 2-3, 3-5, 5-10, 10-20. Less than 50% of the *m/z* in most intervals matched in HMDB and PubChem accurately, with larger ppm intervals leading to more incorrect matches (Figure 1c, Figure S1a-b). For instance, within the 0-1 ppm range, approximately 3000 *m/z* from MoNA and 1000 *m/z* from GNPS were found, but only around 200 and 500 matched in HMDB. Notably, using PubChem reduced matching accuracy significantly. These findings highlight the need for a more comprehensive database and a new pipeline to enhance formula assignment accuracy, minimizing reliance on ppm matching alone.

### SMART Database Construction

Figure 1a illustrates the workflow of Spatial Metabolite Formula Predictor (SMART) approach, highlighting two key features: building a comprehensive database of unique formulas, and machine learning-assisted candidate ranking. Formulas with molecular weight exceeding 1000 and those not following IUPAC nomenclature (Supplementary Table 2) were excluded, leaving 20,154 (9.25%) from HMDB, 346,214 (15.02%) from ChEMBL, and 2,844,720 (41.68%) from PubChem (Figure S1c). Overlaps among these yielded 2,852,349 unique formulas, refined into KnownSet (Figure S1d). Chemical differences between KnownSet formulas produced over one billion shifts, termed as DBEdges (Figure 2a).

**Fig 2.**
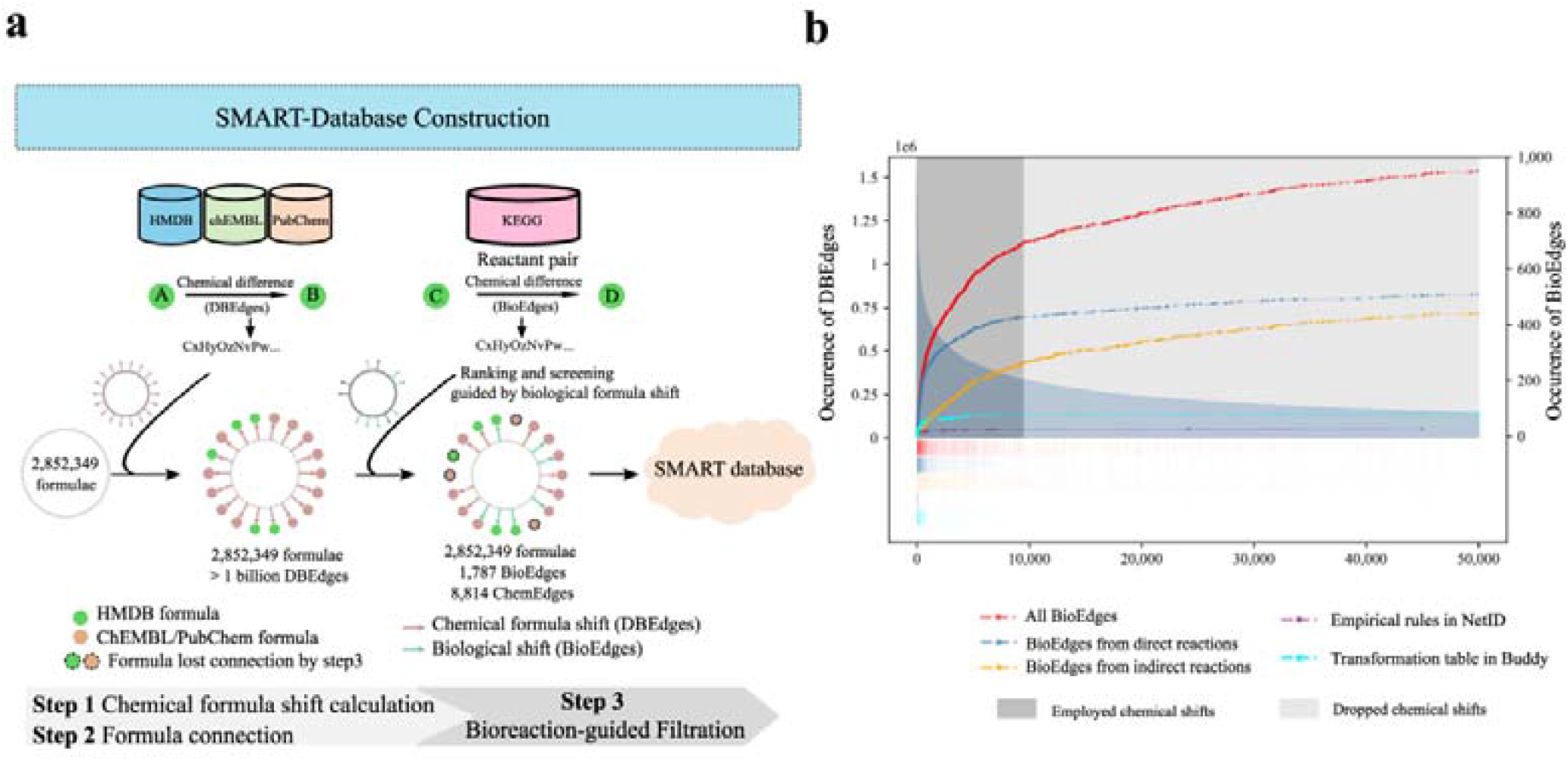
SMART database construction and dataset generation. a. Construction of the SMART database. b. The occurrence of DBEdges, BioEdges with direction reaction shifts or indirect reaction shifts, and the reported reaction rules in NetID and Buddy.

To filter nonsensical connections due to synthetic compounds in PubChem and ChEMBL, we used a biological reaction-guided process. KEGG provided 7,905 reaction pairs: 5,882 direct reaction pairs (DR), and 2,023 indirect reaction pairs (IDR). Here DR represents direct metabolite links, while IDR refers to unresolved R-group or chemical repeats. We calculated the biological formula shifts both from DR and IDR pairs to form BioEdges (Supplementary Table 3). When ranking BioEdges by DBEdges frequency, 510 (73.8%) shifts from DR and 442 (37.3%) from IDR were in the top 50,000 DBEdges (Figure S1e). Using a slope cutoff of 0.001, DBEdges beyond rank 9,508 were excluded (Figure 2b). Consequently, our formula shift database includes 1,787 BioEdges and 8,814 ChemEdges (other DBEdges except BioEdges) (Figure S1f). BioEdges were enriched in C-H-O shifts, while ChemEdges contained more C-H-N-O shifts (Figure S1g). Ultimately, the SMART database interconnected all 2,852,349 formulas via 8,814 ChemEdges and 1,787 BioEdges (Figure 2a).

### Formula Annotation and Evaluation

To annotate measured *m/z* using SMART database, we built a multiple linear regression (MLR) model incorporating four crucial features: Sn (number of nodes linked to each candidate), BioEdges (biological shift connected to each candidate), ChemEdges (chemical shift edges connected to each candidate), and ppm (mass error). The model was trained using NetID dataset of 118 metabolites with *m/z* errors under 3 ppm (Supplementary Table 4). The results indicated that lower ppm errors, and higher Sn, BioEdge and DBEdge yielded more accurate formulas predictions (Figure S2a). We further tested the method on GNPS dataset with 3280 metabolites, showing similar trends (Figure 3a).

**Fig 3.**
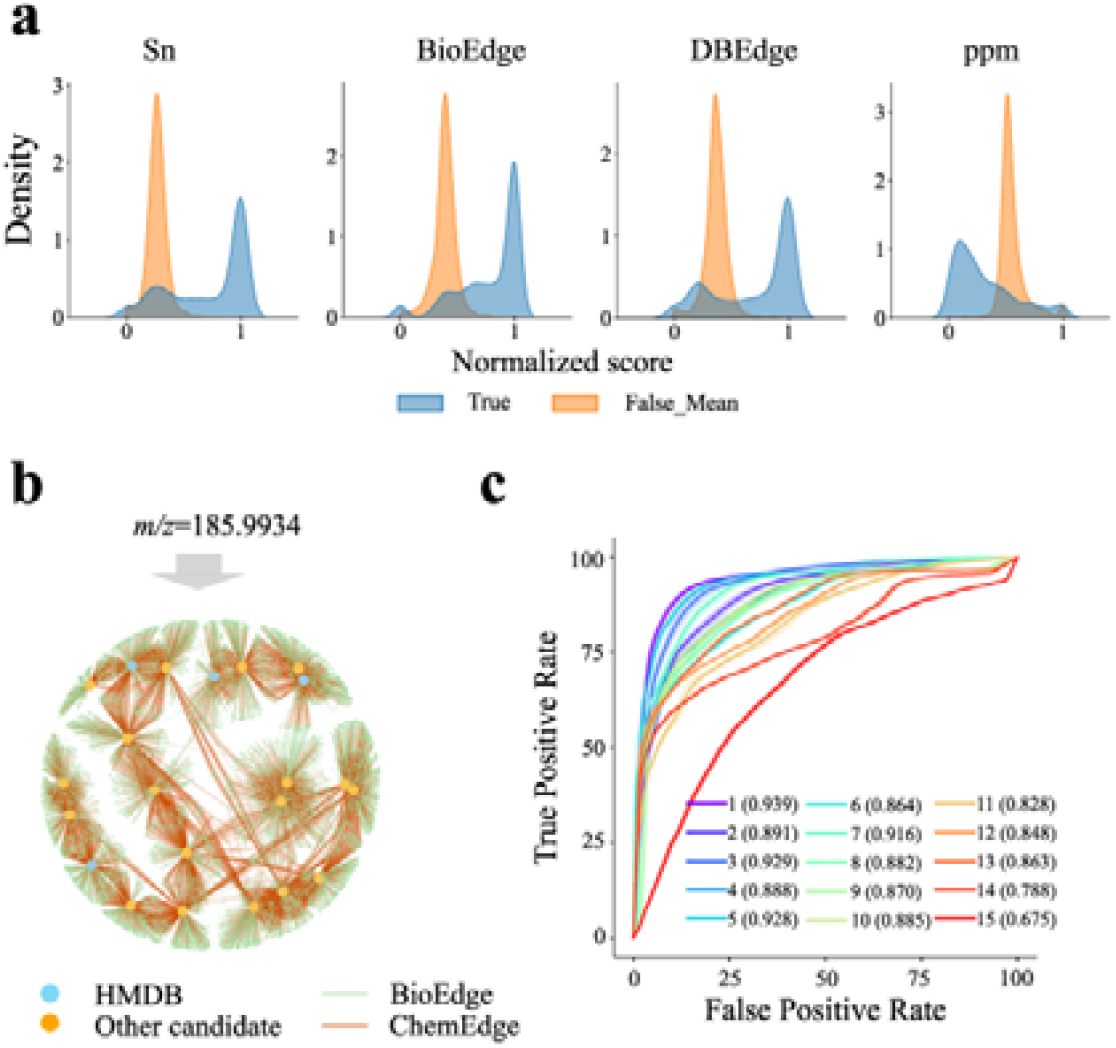
Formula assignment in SMART. a. Feature distribution of formula candidates in the SMART database within a 5 ppm for GNPS dataset. For each *m/z*, features were normalized by dividing the maximum value. The blue color represents the true formula and yellow color represents the mean value of false formulas. b. Formula assignment for *m/z* 185.9934. The network for *m/z* 185.9934 was constructed by matching the *m/z* value with formula candidates within a defined ppm tolerance in SMART database (b). In the network, large circles filled with colors represent candidates from different datasets (HMDB: blue, Other candidates: orange). All these candidates were linked to the formula in SMART database (small green circles) via BioEdges (green line) or ChemEdges (brown line). Four features of formula candidates were extracted and normalized from the network including ppm, Sn, BioEdge, and ChemEdge. Scores were calculated by SMART MLR model. (c).ROC analysis on GNPS datasets using models that randomly combined the four features including: 1, Combined; 2, BioEdge_DBEdge_PPM; 3, Sn_DBEdge_PPM; 4, Sn_BioEdge_PPM; 5, Sn_BioEdge_DBEdge; 6, Sn_BioEdge; 7, Sn_DBEdge; 8, Sn_PPM; 9, BioEdge_DBEdge; 10, BioEdge_PPM; 11, DBEdge_PPM; 12, Sn; 13, BioEdge; 14, DBEdge; 15, PPM. The values in parentheses are the average of AUC values.

To illustrate the functionality of the MRL model works, we consider *m/z* 185.9934 as an example. In a kidney extract sample, *m/z* 185.9934 was identified as 3-phosphoglyceri acid, with an error of 2.6 ppm using LC-MS (Figure S2b and c). SMART linked candidate formulas through edges forming a network that expanded as ppm tolerance increased (Figure 3b). At 5 ppm, 13 candidates, including the correct one (C_3_H_7_O_7_P) were identified (Figure S2b). While some candidates (e.g., C_4_H_11_O_2_PS_2_) offered closer ppm tolerances, the MLR model ranked C_3_H_7_O_7_P highest (Figure S2b), highlighting the limitations of ppm matching alone. To assess the importance of the four features in the MLR model, we performed ROC analysis on both GNPS and MoNA datasets. Results showed that the combination of all four features achieved the highest AUC at 0.939 for GNPS and 0.915 for MoNA, respectively (Figure 3c and Figure S2d). Metric like accuracy, precision, sensitivity and specificity (Figure S2e and f) further confirmed the performance with all features combined.

SMART’s accuracy was further validated against SIRIUS^10^, BUDDY^11^ (without MS/MS information), and ppm matching across four published datasets (American Gut, Tomato, Chagas Diseases and NIST human feces projects)^11^. Our method achieved high Top 1 accuracy rates, with an average of 83.8% across all datasets, and an average Top 3 accuracy rate of 92.4%, surpassing the average performance of SIRUS (71.5%) and BUDDY (80.4%). (Figure 4a, Supplementary Table 5-9). In high ppm tolerance datasets (e.g., GNPS and MoNA), SMART maintained high accuracy (Top1 accuracy of 64.2% and a Top3 accuracy of 79.5%) compared to BUDDY (Top1 accuracy<50%) (Figure 4b, Supplementary Table 10-12). These results confirm SMART’s reliability for formula assignment with only *m/z* value, even without RT and MS/MS.

**Fig 4.**
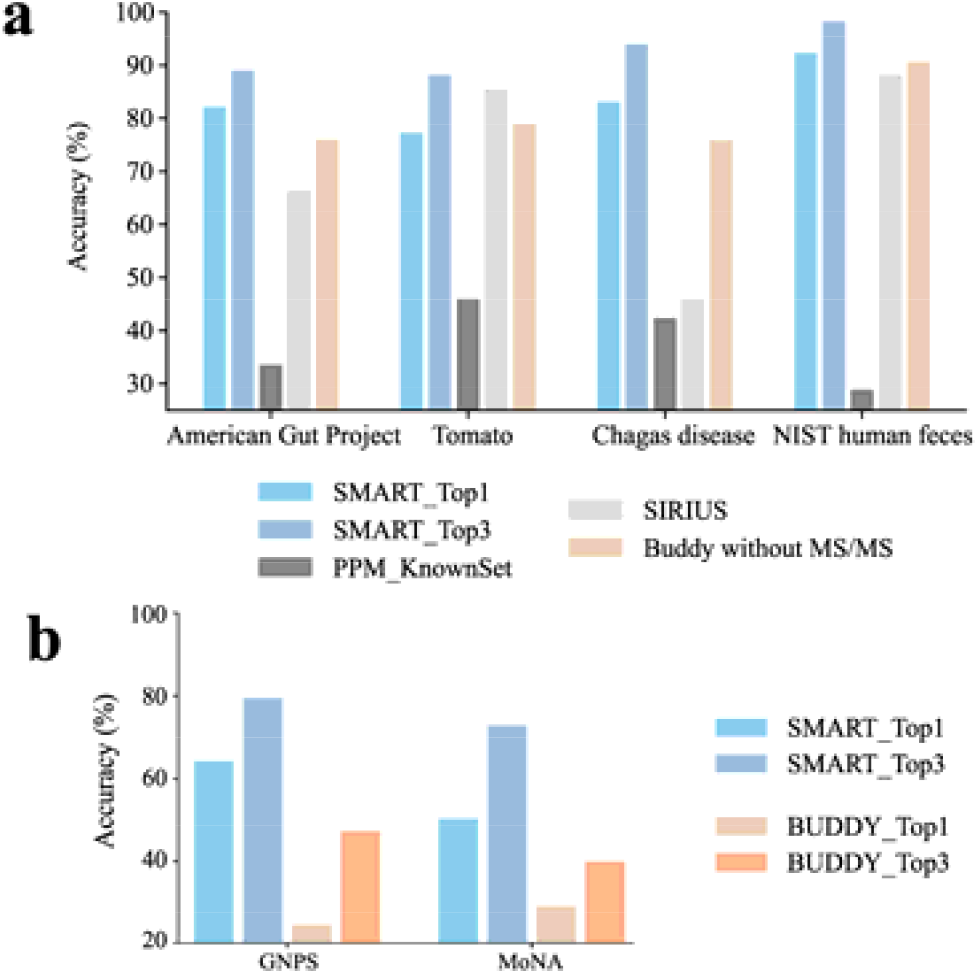
Accuracy evaluation of SMART. a. Top1/Top3 accuracy of SMART compare to other tools for four published datasets (American Gut Project, Tomato, Chagas Diseases and NIST human feces). PPM_KnownSet refers to using ppm matching against the KnownSet database. b. Top 1/Top 3 accuracy of SMART (using 5 ppm) and BUDDY (without MS/MS) against GNPS and MoNA datasets.

### Metabolomics Changes During Mouse Embryo Development

Embryonic development involves complex metabolic reprogramming within each organ, highlighting the need for a detailed study of metabolic regulation throughput embryogenesis^34^. Using MSI-based analysis, we examined mouse embryos at day 14.5 (E14.5) and day 18 (E18) (Figure 5a), detecting 1,357 and 1,431 *m/z* peaks, respectively, with 1,023 shared (Figure 5b). Isotope pattern matching and manual reviewed revealed that over 30% (347 of 1,023) of the shared peaks lacked isotope information due to low signal intensity or overlapping signals (Figure S3a). As a result, tools like pySM, rMSIannotation were insufficient for annotating all *m/z* peaks, whereas SMART successfully annotated 986 *m/z* values (96.4% of the 1,023 peaks).

**Fig 5.**
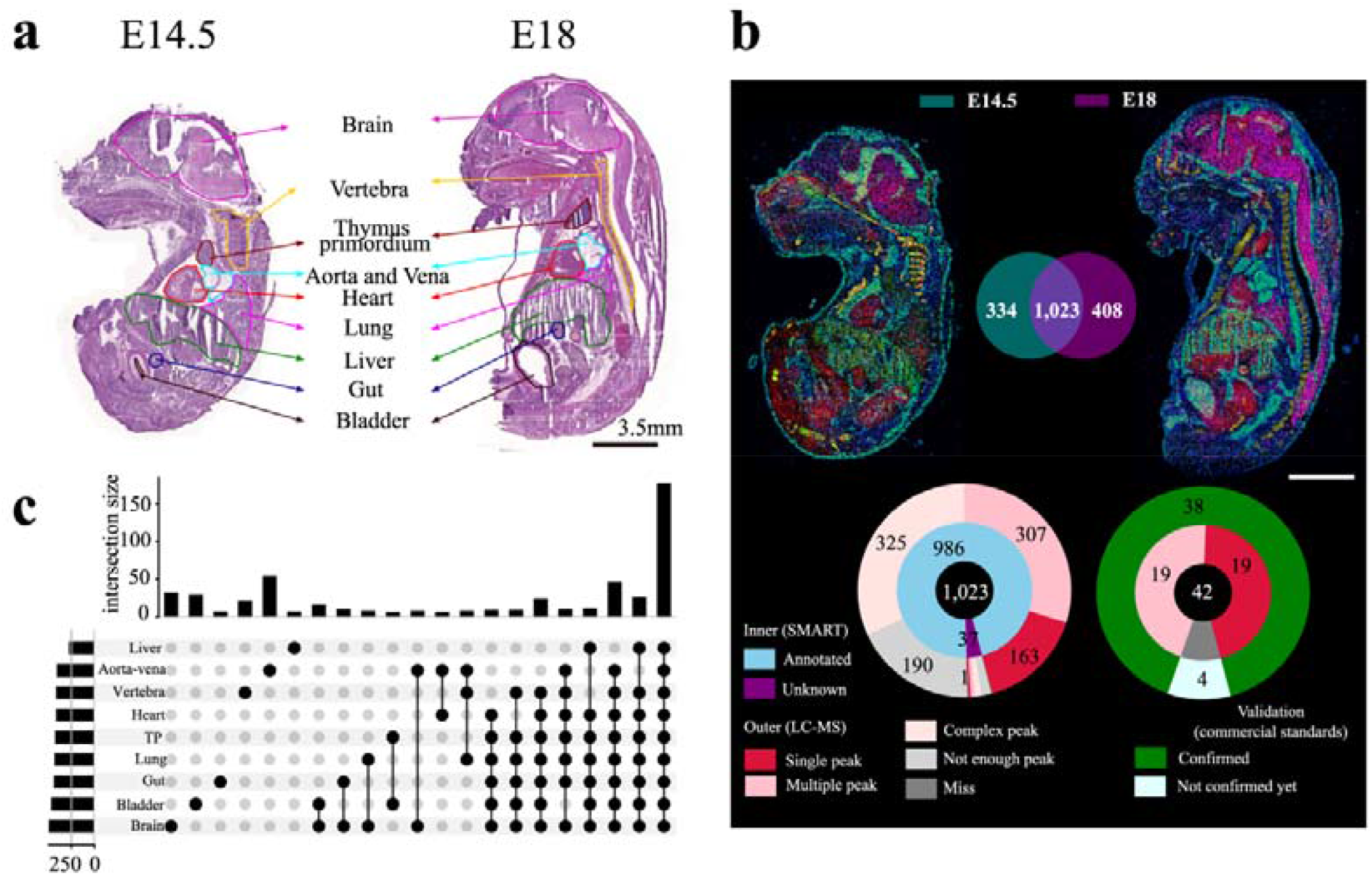
Metabolomics changes during mouse embryo development. a. H&E staining of mouse embryos from E14.5 and E18. Colored outlines designate nine regions of the embryo. b. Statistics of peaks in MSI results for E14.5 and E18 of mouse embryos. Formula assignment and validation was performed for shared 1,023 candidate peaks. The validation results were classified into five types according to the peak complexity (single, multiple, complex, not enough peak and missing). c. An upset plot illustrates the intersection size with more than five candidates across nine organs at E14.5 and E18, with the number of peaks detected in each organ indicated in the left bar.

To confirm the uniqueness of these annotated formulas, we performed LC-MS-based metabolomics on extracted mouse embryo samples. Among the formulas candidates, 163 (21.63%) yielded singular peaks in LC-MS within 5 ppm, indicating their unique identities without interference from isomers (Figure 5b, Figure S3b). Additionally, 311 (34.01%) of these formulas generated multiple peaks in LC-MS, suggesting the presence of multiple metabolites corresponding to a single formula. For instance, C_6_H_12_O_6_ could represent glucose, fructose or inositol. These findings indicate that additional analysis such as MS/MS, derivatization, or ion mobility, are necessary to reliably identify metabolites and differentiate isomers. Of the formulas detected in both MSI and LC-MS, 38 were further validated and identified by matching *m/z* values and retention time with purchased standards (Supplementary Table 13). For instance, C_4_H_9_N_3_O_2_ eluting at 14.49 mins was confirmed as the creatine, and C_6_H_14_N_2_O_2_ eluting at 20.11 min was identified as lysine (Figure S3c-d).

It has been found that glucose levels in the brain are higher than in other tissues, and glycolytic intermediates such as glucose-6-phosphate (G6P) elevates in both the liver and brain after E12^35^. We found high signal intensity of glucose and G6P via examining their spatial distribution in our MSI results (Figure S3e). To further explore metabolic changes during embryonic development from a spatial perspective, we aligned Hematoxylin and Eosin (H&E) images of embryos with MSI data to identify key organs (e.g., brain, gut, heart, liver) (Figure 5a) and reveal organ-specific metabolic profiles The brain had the most assigned formulas, while the liver showed the fewest (Figure S3f). 176 formulas were detected across all organs with varying intensities, and others were specific to particular organs, such as 31 formulas in the brain and 6 in the liver (Figure 5c). Among all organs, the liver displayed the greatest number of formulas with signal levels rising at E18 compared to E14.5 (Figure S3g). Conversely, brain signals decreased significantly (Figure S3g). For example, C_10_H_15_N_3_O_5_, the top 1 candidate for *m/z* 257.1021, was enriched in the brain, liver and thymus primordium at E18 (Figure S3h). LC-MS analysis confirmed it as 5-methylcytidine, which plays a critical role in embryonic stem cell self-renewal and differentiation^36^. Similarly, C_7_H_15_NO_3_ (carnitine) was enriched in liver, bladder, and lung at E18 (Figure S3i) that facilitating fatty acid beta-oxidation and thereby supporting embryonic development^37^. Thus, SMART can accurately annotate formulas solely based on *m/z* information, enhancing MSI metabolite identification and related biological studies.

## Discussion

Organs are composed of distinct spatial regions and cells types, varying in proximity to blood vessel for nutrient acquisition. Variations in transporters and enzymes expression further amplify metabolic differences, even among neighboring cells. Current MSI techniques such as MALDI or DESI generate peaks with *m/z* information but lack robust MS/MS fragmentation due to the low sensitivity and miss RT because of the absence of pre-separation steps. Therefore, traditional pipelines that rely on RT or MS/MS data are unsuitable for MSI data analysis. Typically, *m/z* values are matched with metabolite databases based on natural abundance, but mass spectrometry accuracy, ppm variation and machine sensitivity limit these approaches to 35% accuracy established databases (Figure 4a). While high abundance metabolites can sometimes be annotated, a substantial gap remains in annotating metabolites with precision.

The development of SMART represents advancement in spatially-resolved metabolomics, addressing the challenges of formula annotation in MSI. Its extensive database and multiple linear regression model enable accurate annotation, as demonstrated by its annotation of 986 formulas in developing mouse embryos. ChemEdges and BioEdges offer insights into possible formula extension. Beyond spatially-resolved analysis, SMART also shows great promise for application in single cell metabolomics, which similarly lacks RT information and face challenges in obtaining robust MS/MS spectra. Moreover, SMART has potential to predict the fragment formula in MS/MS, which is crucial for high-throughput metabolite structure inference.

While SMART has demonstrated considerable progress, there are still opportunities for further improvement. A single formula can represent multiple metabolites (e.g., glucose, fructose, inositol all give formula as C_6_H_12_O_6_), and issues like adducts and in-source fragmentation complicate detection. Parallel LC-MS runs often validate only a fraction of MSI results. In our study, 38 validations of formulas in mouse embryo MSI covered less than 10% compared to LC-MS runs using an in-house standard library. Therefore, enhancements, such as incorporating additional metabolite data, refining machine learning algorithms, on-tissue derivatization, or ion-mobility mass spectrometry, are necessary to improve annotation accuracy.

In conclusion, SMART’s systematic annotation capability makes it as a transformative tool in the field of spatially-resolved metabolomics. Its comprehensive database, well-established multiple linear regression model and high accuracy make it indispensable for researchers to unravel the complexities of tissue-specific metabolism. By providing a detailed spatial map of metabolites, SMART can advance our understanding of metabolic processes in two-dimension.

## Methods

### Generation of KnownSet and the datasets for evaluation

To construct the KnownSet database of chemical formulas, data were sourced from three databases: HMDB, ChEMBL, PubChem. The following three filtration steps were applied: (i) chemical structures with charges and a molecular weight exceeding 1000 were excluded from the dataset (ii) formulas containing atoms not recognized by IUPAC were excluded. (iii) duplicates were merged, ensuring that each formula was unique across the three database.

The NetID dataset was manually curated and downloaded from supplemental table 2 of the NetID^9^. All artifacts and isotopes were removed during processing. For the GNPS and MoNA datasets, experimental spectra in json format were downloaded from their official websites (GNPS, https://gnps.ucsd.edu/ProteoSAFe/static/gnps-splash.jsp; MoNA, https://mona.fiehnlab.ucdavis.edu/). Only candidates with *m/z* values below 1000 and a ppm error less than 20 were retained. Other four datasets with annotation results, American gut, Tomato, Chagas Diseases, and NIST human feces datasets, were downloaded from supplemental table 4-7 of BUDDY^11^.

### Definition of ppm intervals in evaluation datasets

For each formula in the GNPS, and MoNA datasets, the ppm error (P) was calculated based on the theoretical values (M_0_) and measured *m/z* values (M_c_) according to the following formula: abs(M_0_ - M_c_)/M_0_*1e6, where abs referred to absolute value function. For ppm matching evaluation, each dataset was divided into six intervals based on ppm error, including 0-1, 1-2, 2-3, 3-5, 5-10, 10-20. For the SMART MLR model evaluation, a cumulative intervals was defined with ppm error located in P<=5.

### DBEdges formula shift calculation

The chemical difference, named as formula shift, was calculated between two seed formulas based on differences in their atomic composition. An illegal shift was defined when both positive and negative atom counts exist simultaneously, which were excluded. The remaining formula shifts were then tallied and ranked according to their frequency of occurrence in KnownSet database. To maintain a stringent cutoff, any formula shift occurring fewer than 5,000 times was excluded from consideration. These filtered shifts exceeding 1 billion are named as DBEdges.

### BioEdges formula shift filtration from biological reaction database

Bio-reactions pairs were analyzed from KEGG REACTION databases. Initially, 7,905 reactant pairs with 7,042 compounds were extracted to generate 1,699 reaction classes. Some compound formulas contain undetermined functional group represented by the symbol R, or lacked a specified number of functional groups, represented by the symbol N. We included all possible formulas extended with different R and N, while excluding any whose molecular weight exceeds 1000 (Supplementary Table 3). As a result, the number of reactant pairs increased to 9,691. Then all reaction pairs were categorized into two groups: (i) formula pairs without R and N symbols, referred to direct reaction pairs (DR) and (ii) formula pairs involving extened formulas with different R or N symbols, referred to indirect reaction pairs (IDR). Formula shifts were then calculated and illegal pairs from each group were excluded only if an illegal shift was present. Formula shifts from group (i) were named as direct reactions shifts and were used to determine the DBEdges threshold in the following paragraph. Formula shifts from group (ii) were named as indirect reaction shift and were used solely for formula connection. All formula shifts are referred to as BioEdges. We computed the cumulative occurrence of direct reaction shifts based on the rank list in DBEdges. By determining of occurrence growth rate versus its ranking of shift, we choose a cutoff slope of less than 0.001 to remove all subsequent DBEdges. Eventually, our final shifts database composed of DBEdges preceding this cutoff and all BioEdges.

### Feature definition and multiple linear regression model (MLR model) construction in SMART

For each *m/z* value, all candidate formulas will be retrieved from formula database with a defined ppm error (5 ppm, in this work). All shifts linking formulas in the SMART database and candidate formulas will also be obtained. These nodes (formulas linking to candidates) and edges (formula shifts) will be used for feature calculation including Sn, BioEdge, DBEdge, and ppm.

For each candidate, Sn is defined as the sum of the linked nodes, represented by the following formula: Sn = ∑ kN_k_, where k ∈ (1,2,3) denotes the number of databases containing the nodes and N_k_ represents the number of linked nodes for each k. The BioEdge feature stands for the number of BioEdges linked to the candidate and DBEdge indicates the number of other DBEdges linked to the candidate. ppm refers to the ppm error between the candidate (M_c_) and input *m/z* value (M_0_) according to the following formula: abs(M_0_ - M_c_)/M_0_*1e6. For each *m/z* value, each feature of a candidate was normalized by dividing it by the maximum value of that feature in all candidates.

To build the SMART model, normalized features were calculated for all the 118 metabolites in NetID dataset. Since each reference metabolite could generate one true formula and many false formulas, the normalized features between true and false formulas are imbalance. To address this phenomanon, an undersampling strategy was employed using RandomUnderSampler, a module from third-party python package called imlearn. A multiple linear regression model was built based on the 118 metabolites with 4 features, and the equation was obtained as: Y=Xβ*+*ε, where X=(Sn, BioEdge, DBEdge, ppm), β*=*(1.6025, 1.5975, -2.2289, -0.2211), ε*=*0.1074. Meanwhile, models based on single features were also constructed to compare the integration of 4 features.

### Evaluation of MLR model construction in SMART

To evaluate the performance of models constructed from NetID dataset, a large collection from the GNPS dataset was used, which contains 25 times more metabolites. Prediction was performed for all metabolites in GNPS dataset. To calculate evaluation attributes such as the AUC scores, the undersampling process was performed 500 times for each model. The accuracy of MLR model was evaluated by ranking the formulas according to the values generated by the model, considering the Top 1/Top 3 predicted formulas. All analysis was performed using the scikit-learn python package. Since only the *m/z* lists from GNPS and MoNA dataset were used with no spectra provided, the evaluation of Buddy^11^ was performed under its ‘mz_to_formula’ function from the python package msbuddy (https://github.com/HuanLab/BUDDY).

### IACUC statement and animal models

All animal studies were approved by the Institute of Basic Medical Sciences (IBMS)/Peking Union Medical College (PUMC) Animal Care and Use Committee (ACUC-A01-2022-003). C57BL/6J pregnant mice were purchased (Vital River, Beijing) and allowed two days of acclimation to the animal facilities before experiment. Mice were housed on a standard light cycle (from 8:00 to 20:00) and fed with a standard rodent chow (Jiangsu-Xietong, XT101FZ-002).

### Statistics and reprehensibility

All statistical analyses were conducted in Python (3.8). Method evaluation and data analyses were performed computationally and can be reproduced by preprocessing pipelines.

## Code availability

SMART is written in the Python language. Source codes with a small formula database extended from HMDB can be freely downloaded from https://github.com/bioinfo-ibms-pumc/SMART. For the entire database construction, please ask author for further help.

## Author contributions statement

Y.C., L. Y. and L. W. conceived the project. Y. C. developed the method and wrote the SMART code. S. L. performed mass spectrometry imaging and LC-MS experiment. S. L. and Z. L. performed the mice experiments. Y. C., Z. J., X. P., J. Z. analyzed the data and draw the figures. Y. C., L. Y. and L. W. wrote the manuscript. All authors discussed the results and commented on the manuscript.

## Acknowledgments

This work was supported by the Beijing Natural Science Foundation (Z240014, L. W.), the Fundamental Research Funds for the Central Universities (3332023039, J. Z.) the Non-profit Central Research Institute Fund of the Chinese Academy of Medical Sciences (2021-RC350-008, L. W.), National Key R&D Program of China (2022YFA1106300, L. W.), State Key Laboratory Special Fund (2060204, L. W.), National Science Foundation of China Grants (32271354, L. Y.), Talent Plan of Shanghai Branch-Chinese Academy of Sciences (CASSHB-QNPD-2023-011, L. Y.), National Natural Science Foundation of China (92353302, J. Z.). We acknowledge the use of High-performance Computing Platform at the Center for Bioinformatics, Institute of Basic Medical Sciences, Chinese Academy of Medical Sciences, School of Basic Medicine Peking Union Medical College. We also thank to Prof. Jia Yu, Prof. Xiaoyue Wang and all members of the Wang laboratories for the discussion.

## Competing Interest Statement

The authors declare no conflicts of interest.

## Notes

### Competing Interest Statement

The authors have declared no competing interest.

